# Biological Innocuousness of a Device developed to select the Optimum Spermatozoa to be used in the Treatment of Infertility

**DOI:** 10.1101/2021.06.23.449593

**Authors:** Marisa A Cubilla, Héctor A Guidobaldi, Laura C Giojalas

## Abstract

The sperm selection assay (SSA) is a method based on chemotaxis to obtain spermatozoa at the optimum physiological state to successfully fertilize the egg. It consists of a device made of acrylic and an attractant solution which includes progesterone. We evaluate potential cytotoxicity interactions by means of Neutral Red uptake, the MTT and colony formation assays, according to ISO normative. Here we showed that even stressing the conditions of the assays, the SSA device alone or together with the progesterone solutions employed, showed to be innocuous for the cells. Suggesting that SSA could be incorporated into the ART procedures.

## Introduction

In our lab, we developed a new device, and the associated method of use, to select spermatozoa at the optimum physiological state to successfully fertilize the egg (e.g. spermatozoa capacitated with low DNA fragmentation and oxidative stress ^[1]^. Thus, the sperm selection assay (SSA; Argentine patent # 070776, USA patent # 8993310, European patent # 2403933, and Japanese patent # 5816101) is based on the ability of capacitated spermatozoa to orient their movement by following a concentration gradient of an attractant molecule, navigation mechanism called chemotaxis ^[2]^. The attractant molecule used in the device was progesterone (steroid secreted by the oocyte surrounding cells by the time of ovulation) which was observed to chemotactically attract capacitated spermatozoa at very low doses ^[3–9]^. The device is made of acrylic and consists of two vertical wells which are connected at the bottom by a transversal tube. Spermatozoa are placed in one well and the attractant solution in the other one, whereas the attractant gradient is formed inside the connecting tube. The device is provided with a hermetic closing system which prevents the liquid flow between the connected wells.

Assisted Reproduction Technology (ART) has been applied to treat infertility during the last 40 years; however, its efficiency is still relatively low (~ 30%), whereas sperm quality is relevant for the ART success. Then, the use of the sperm subpopulation selected by progesterone in the SSA device may improve the outcome of ART; however, cytotoxicity of the device and the attractant solution must be tested before routinely incorporating it to the ART procedure. Then, we first selected the appropriate cytotoxicity assay to test the innocuousness of the SSA device and solutions, and afterwards, we set them up according to the SSA experimental conditions.

In vitro cytotoxicity tests are used to evaluate devices and materials for medical and biological purposes. In recent years, several tests and criteria have been developed to evaluate cytotoxicity in cells which are validated by the standard ISO 10993-5 ^[10]^, whereas the most recommended methods for biomedical applications are the MTT, the neutral red uptake and Colony formation assay, all of them indicating cell viability. The first one evaluates cell viability as the metabolic reduction of MTT (3-(4,5-dimetiltiazol-2-il)-2,5-diphenyltetrazoliumbromid) in mitochondria, which is proportional to the number of viable cells in the exponential growth phase ^[11]^. The neutral red uptake is based on the ability of viable cells to incorporate the supravital dye neutral red. This weakly cationic dye penetrates cell membranes by non-ionic passive diffusion and concentrates in the lysosomes ^[12,13]^. Cell survival is determined after incubation with neutral red by colourimetric analysis of this dye extracted from the lysosomes ^[14,15]^. Colony formation evaluates cell viability as the ability of a single cell to proliferate forming a large colony ^[16,17]^. Here we showed the innocuousness of the SSA by means of selected cytotoxicity methods recommended by the ISO normative.

## Materials and Methods

### Cell preparation

We used the cell line L-929 (ATCC-CCL-1) recommended by “ISO 10993-5” ^[10]^ to test cytotoxicity with high sensitivity. The cell line was maintained in Eagle’s Minimum Essential Medium (MEM; Gibco-BRL, Gaithersburg, MD, USA) supplemented with 10% fetal bovine serum (FBS; Gibco-BRL, Gaithersburg, MD, USA), 100 units/mL penicillin G-streptomycin and 4 mM L-Glutamine (293 mg/mL). Further cells incubation was carried out according to each cytotoxicity assay.

### Extract solution preparation

The SSA devices were cleaned with 1N HCl for 24 h, and then gently washed with distilled water, and dried in an incubator at 40°C. Then, the devices were sterilized in a microwave oven at 900W for 3 min. The extract solution was prepared by loading the SSA wells with MEM (standard culture medium for L929 cell lines) or HTF (Human Tubal Fluid medium, used for sperm culture) with or without progesterone for 24 h. Then, the extract solution samples were recovered and kept at −80°C until the moment of use. Additionally, MEM and HTF culture medium were incubated for 24 h, in polypropylene tubes as a control of an innocuous material for cell growth (Blank solution). It is known that PBS does not support cell survival for the long term because it is a simple salt solution without essential cell survival elements; hence, it was used to dilute the extract solutions with the purpose to induce stressing suboptimal cell growth to enhance the cytotoxicity effects, if any. Solutions preparation are summarized in table 1. Then, each of the tested solutions was added to the cells and further incubated for 24, 36 or 48 h, in an atmosphere containing 5% CO_2_ in the air, at 37 °C. At the end of the incubation time, the different viability tests were performed, as described below.

**Table 1.** Description of Extract Solutions Preparation

|  |  |
| --- | --- |
| **MEM** | Eagle minimum Essential Medium with 10% fetal bovine serum (FBS), 100 units/ml penicillin G-streptomycin and 4mM L-Glutamine. |
| **HTF** | Human Tubal Fluid Culture Media used in Sperm Selection Assay (SSA). |
| **MEM<sub>SSA</sub>** | MEM culture medium after 24 h of incubation with SSA devices. |
| **HTF<sub>SSA</sub>** | HTF culture medium after 24 h of incubation with SSA devices. |
| **HTF<sub>SSA+P</sub>** | HTF culture medium with Progesterone after 24 h of incubation with SSA devices. |
| **BLANK** | Culture Medium (MEM) incubated in polypropylene tubes (Falcon Becton Dickinson). |
| **PBS** | Phosphate buffer solution pH 7.4. |

### MTT assay

Cells were incubated in a 96-well tissue culture plate at 37°C in an atmosphere containing 5% CO_2_ in air until a semi-confluent monolayer was achieved. Then, MEM was replaced by 200 μL of extract solution and further incubated for 24, 36 or 48 h. After that, the extract solution was replaced by 50 μL of MTT solution, followed by a 3 h incubation. The MTT solution was removed, and 100 μL of Isopropanol was added. Then, the level of reduced MTT was determined by measuring the optical density (OD) of the solution in a microplate reader (Elx800 Absorbance Microplate Reader Imager software GEN5, Biotek Instruments, USA) at λ = 570 nm. In all treatments, the percentage of viability was calculated by dividing the OD of the extract solution by the OD of the blank solution per 100.

### Neutral red uptake assay

Cells were incubated in a 96-well tissue culture plate at 37°C in an atmosphere containing 5% CO_2_ in air, until a semi-confluent monolayer was achieved. Then, MEM was replaced by 200 μL of extract solution and further incubated for 24, 36 or 48 h. After that, the extract solution was discarded, and cells were washed with 100 μL PBS. Then, 100 μL of Neutral Red (NR) solution was added to each well. After 4 h of incubation, the NR was removed and cells were washed with 100 μL PBS, and 150 μL of NR desorbing fixative solution, composed of 50% ethanol:1% glacial acetic acid, was added. The uptake of NR was determined by measuring the OD of the solution in a microplate reader (Elx800 Absorbance Microplate Reader; Imager software GEN5, Biotek Instruments, USA) at λ = 540 nm. In all treatments, the percentage of viability was calculated by dividing the OD of the extract solution (minus the OD of PBS) by the OD of the blank solution (minus the OD of PBS) per 100.

### Colony formation assay

The cell suspension containing 6×10^4^ cells/well were seeded into six-well plates and maintained at 37°C in an atmosphere containing 5% CO_2_ in air for 24 h. Then, each well was washed with PBS and 2000 μL of extract solution were added, incubating the cells for 7 days. Every 24 h the plates were observed under a phase contrast microscope (Nikon Ti-S/L100, USA) and the total number of colonies per well was determined. Colonies consisting of 50 or more cells in each well ^[10]^.

### Statistical Analysis

Extract solutions and incubation time on cell viability were evaluated using statistical software GraphPad Prism 5.01 (GraphPad Prism version 7.00 for Windows, GraphPad Software, La Jolla California USA, www.graphpad.com”). We used XY analyses of column Statistics for calculating descriptive statistics and differences between extract solutions were determined by means of a one-way ANOVA, and a posteriori Tukey test considering statistically significant differences at *p* ≤ 0.05. The correspondent dataset are available at https://data.mendeley.com/datasets/6ygpz8562x/draft?a=13579ccf-a612-44fa-af06-532b4279b8ae

## Results

In relation to the MTT assay, all MEM dilutions tested showed viability higher than 70% (the lower limit imposed by the norm to be considered as no toxic effect), at least until 48 h of incubation, in contrast to the survival of cells under the PBS solution (Fig. 1A). In the case of the HTF, most of the dilutions preserve cell viability along time over the threshold of 70% (in contrast to PBS treatment), except for the dilution of 25% HTF:75% PBS after 48h of incubation, a condition that showed a slight decrease below the threshold (Fig. 1B). All dilutions of the MEM_SSA_ extract showed a decrease in cell viability after 24 h of incubation, which was later recovered until 48 h (Fig. 1C). However, only the 25% MEM_SSA_:75% PBS dilution remained below the threshold at least until 48 h. When cells were exposed to the HTF_SSA_ extract a decrease in cell viability was observed in all dilutions after 24 h of incubation, which was followed by a recovery in all treatments (Fig. 1D). Cell viability under the HTF_SSA+P_ was over the threshold along the incubation time, whereas only the higher stressing condition (25% HTF_SSA+P_:75 %PBS) showed a value a slightly lower than the 70% viability threshold, at least until 48 h (Fig. 1E). When cell viability was determined by the NR assay, a similar behaviour to MTT outcome was observed (Fig. 2).

**Figure 1.**
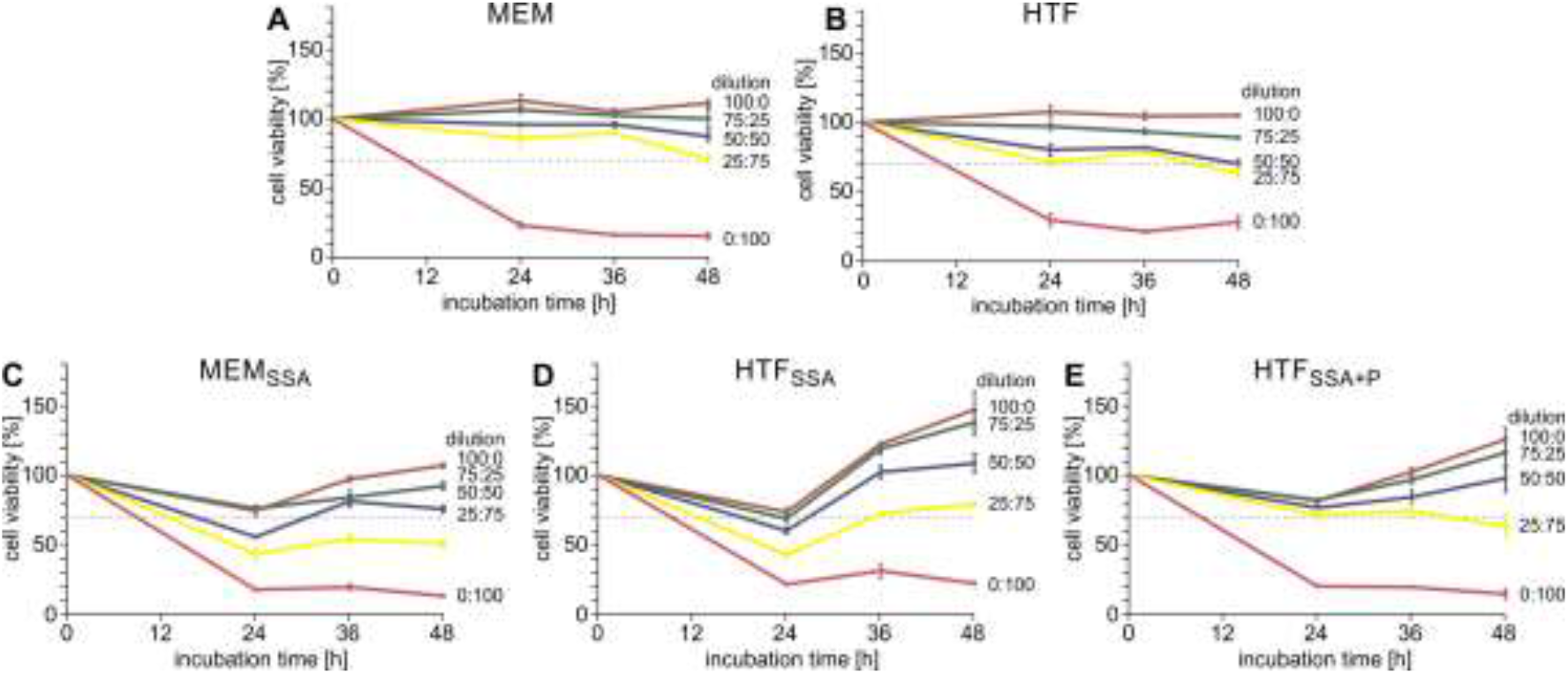
Cell viability measured by MTT Assay. Cells were incubated with different dilutions of extract solutions: A) MEM, B) HTF, C) MEM_SSA_, D) HTF_SSA_ and E) HTF_SSA+P_ and the viability was evaluated by MTT assay at different incubation times. Dash line at 70% of cell viability, indicates the limit imposed by the norm to consider cytotoxic effects. Dilution indicates the proportion of extract solution to PBS.

**Figure 2.**
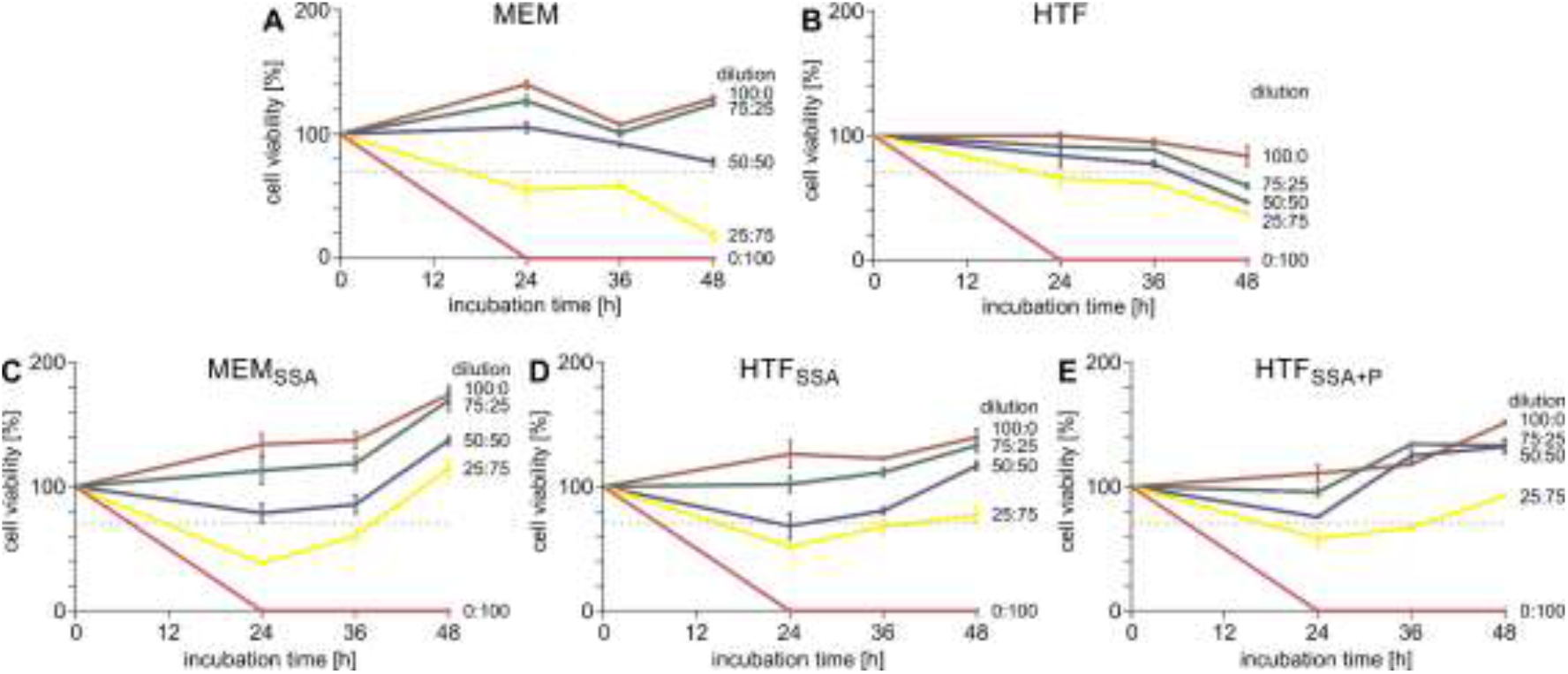
Cell viability measured by Neutral Red Assay. Cells were incubated with different dilutions of extract solutions: A) MEM, B) HTF, C) MEM_SSA_, D) HTF_SSA_ and E) HTF_SSA+P_ and the viability was evaluated by Neutral Red assay at different incubation times. Dash line at 70% of cell viability, indicates the limit imposed by the norm to consider cytotoxic effects. Dilution indicates the proportion of extract solution to PBS.

As observed in figure 3, all the extract solutions evaluated promote the growth of cells colonies, as compared to the respective control. The results suggest that the SSA components (device and HTF with P) did not affect cell viability.

**Figure 3.**
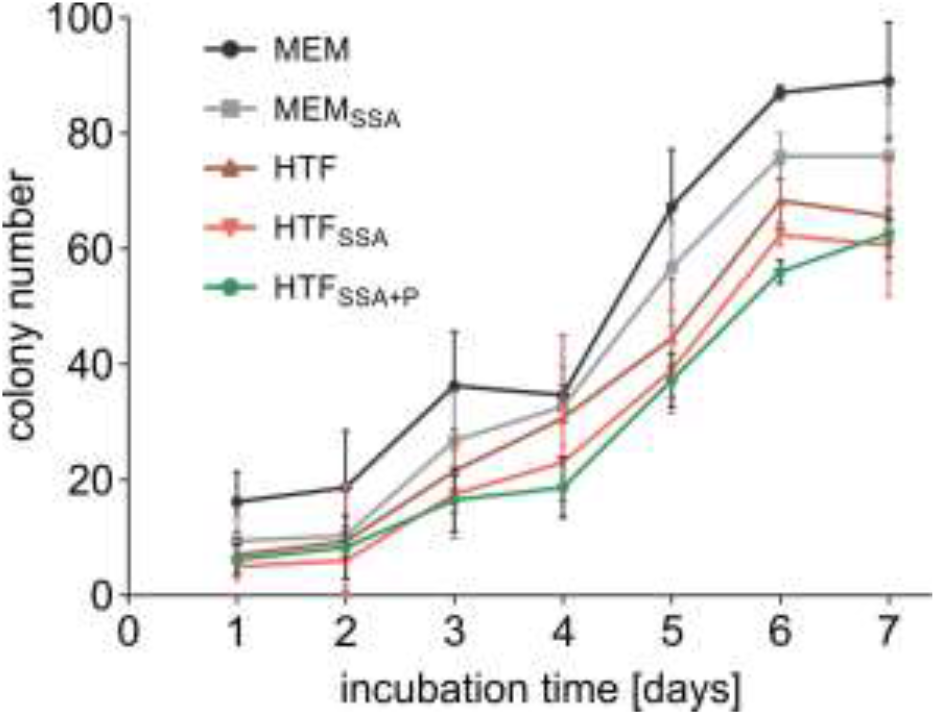
Cell colonies growth measured by Colony Assay. Cells were incubated with different dilutions of extract solutions: A) MEM, B) MEM_SSA_, C), HTF, D) HTF_SSA_ and E) HTF_SSA+P_ and the cell colonies number was evaluated at different incubation days.

## Discussion

Cytotoxicity evaluation is a primary requirement for the use of medical devices with patients, whereas the main cytotoxicity assays are based on cell survival and growth under in vitro incubation with the device extract solution as recommended by the International Standard ISO 10993-5 ^[10]^. Here we used three different assays (MTT, NR and colony assay), which evaluate the potential cytotoxicity of the SSA components (the acrylic device, the specific culture medium and progesterone) according to different biological mechanisms.

When the extract solutions were prepared with MEM supplemented with serum, both polar and no-polar substances are expected to be extracted ^[10]^. In the case of HTF, which is also a salt solution supplemented with synthetic serum, chemicals could (or could not) also be extracted from the device upon HTF interaction, with or without progesterone.

In general, for most of the tested extracts, the MMT and NR assays showed cell survival above the 70% threshold; however, under the most cell stressing extract dilutions, after 24 h of cell culture, a slight decreased was observed. Conversely, such decay is recovered for most experimental conditions at 48 h. The colony formation assay, showed that the cell growth in any kind of extract along time, suggesting that the SSA components did not affect it. In all event, the HTF extracts were slightly lower than those of MEM, an observation that was not associated with a toxic effect since the HTF composition is not optimum for the cell line growth. It is worth to note that even 24 h of incubation is ~70 folds higher than the 20 min in which the cells are exposed to the SSA.

Here we showed by mean of three independent normalized assays, that the SSA seems to be innocuous for cell survival, at least in the conditions tested in this study.

## Acknowledgements

MAC, HAG and LCG are researchers from the Consejo de Investigaciones Científicas y Técnicas (CONICET). The study received financial support from the Ministerio de Ciencia y Tecnología (FONTEC 2010; Córdoba government) and Ministerio de Ciencia, Tecnología e Innovación Productiva (FONCYT – Startup – 2010–2070; Argentina government).

